# Evidence of play behavior in captive California Two-Spot Octopuses, *Octopus bimaculoides*

**DOI:** 10.1101/2024.08.23.609397

**Authors:** Katarina Jarmoluk, Galit Pelled

**Author notes:** Corresponding author (GP).

## Abstract

Play is considered to be an essential part of development that supports learning, memory, and the development of flexible behavioral strategies. It may also serve as an informative factor in assessing an animal’s welfare state and in improving care and husbandry practices. An increasing number of non-mammalian species have been discovered to engage in play behavior, including several cephalopod species. Here, we characterized play behavior in wild-caught, laboratory-housed California Two-Spot Octopuses, *Octopus bimaculoides*, a species with growing relevance as a model in biomedical research, with the goal of establishing a behavioral repertoire and encouraging further research into the behavior and welfare of this species.

## Introduction

Play is an essential part of animal development, as it is known to engage neurobiological mechanisms that support learning, memory, and the development of flexible behavioral strategies. It is also important in adulthood, and both human and animal research suggests that playful behavior and internal motivation are linked to mental health [1].

When play behavior first started growing in popularity as a field of scientific study, the only non-human animals thought to be capable of it were other mammals and possibly birds [2, 3], but over the past few decades there has been growing appreciation of the importance of play in development, socialization and communication, as well as an increasing amount of evidence showing that play behavior is also found in reptiles [4, 5], fish [5-7], and even insects [8].

Octopods are soft-bodied marine mollusks of over 300 known species within the class Cephalopoda, containing two major groups: the finless incirrate octopuses and the finned cirrate octopuses. Within the taxon of incirrate octopuses is the family Octopodidae, containing the majority of octopods at over 200 species. Octopodidae are benthic creatures and are characterized by their eight arms, each possessing one to two rows of suckers [9]. The majority of an octopus’ neurons are located within the arms [10, 11], giving it a decentralized nervous system. They are also capable of changing the color and texture of their skin, which they commonly use as a method of camouflage or startle display [12-14].

With their large brains, unique nervous system, and wide behavioral repertoire, octopuses are ideal models in a broad variety of fields and subjects, including aquaculture, neurobiology, climate change, and more [15]. In particular, they are becoming increasingly common as a model in biomedical and neuroscience fields, as well as in robotics [16-19].

The discussion of play in octopuses is not a new occurrence. Play or play-like behaviors have been observed and recorded in two species: the common octopus, *Octopus vulgaris* [20], and the Giant Pacific Octopuses, *Octopus dofleini* [21]; it has been anecdotally described in the Caribbean Reef Octopus, *Octopus briareus* [22]; and briefly mentioned in regards to the California Two-Spot Octopus, *Octopus bimaculoides* [23]. *O. bimaculoides* (Pickford and McConnaughey, 1949) is a species endemic to the coasts of California and Mexico’s Baja Peninsula. This species is one of the most common species of “pet” octopuses and is also growing in popularity as a model for biomedical research [24]. Our current work marks the first time, to our knowledge, that play behavior has been recorded and quantified in *O. bimaculoides*, and that video and photo evidence have been used to characterize play in any octopus species.

Understanding octopus play may be valuable for assessing an animal’s welfare state. It is well-understood that an animal’s welfare state affects the quality of scientific research. An animal’s body is affected by its state of mind [25] and inadequate welfare can cause abnormal behavior, physiology, and immunology [26]. The most important measure for ensuring the protection of cephalopod welfare in both laboratory and field settings is the refinement of methods of maintenance, care, and culture [27, 28]. According to Sykes & Gestal [29], “The practice of good (positive) or bad (negative) welfare in [cephalopod] research, maintenance, rearing or culture conditions will determine the existence of pathologies.”

Existing research shows that stress negatively affects both *O. bimaculoides* physiology and behavior [30, 31] and may be related to stereotypic behaviors such as autophagy [32]. For captive cephalopods, enrichment is an important management tool that can be used to overcome the inherent stress of being held in captivity [33]. Ahloy-Dallaire et. al. [34] suggested that there is a consistent relationship between animal welfare and most types of play. Evidence suggests that positive affect increases play, while negative affect can significantly suppress many types of play in both human and non-human animals. Since there have been very few deliberate records and measurements of octopus play behavior, the association between play and welfare in the Octopodidae family, and in *O. bimaculoides* specifically, remains to be determined. As it grows in popularity as a research model, the importance of establishing a detailed behavioral repertoire for *O. bimaculoides* increases as well.

In this paper, the behavior of *O. bimaculoides* in a lab setting was analyzed in order to characterize play behavior within this species and to demonstrate how a deeper understanding of play in captive octopuses may aid in improving and maintaining octopus welfare. Although the presented findings are preliminary in nature, the authors hope that the presence of such findings may encourage further research into the behavior and welfare of this increasingly biomedically relevant species.

## Methods

All procedures were approved by the Institutional Animal Care and Use Committee at Michigan State University. Nine adult *O. bimaculoides* were obtained off the coast of southern California by Aquatic Research Consultants then shipped overnight to the lab at Michigan State University. Octopuses were housed separately in the lab for an average of 93 ± 43 days. Welfare was regularly assessed using a slightly modified version of the Giant Pacific Octopus health and welfare assessment tool developed by Holst and Miller-Morgan [35].

Each octopus was housed individually in a 40-gallon tank with a separate sump compartment for filtration. Each tank system was isolated from the others to prevent possible water contamination. Water parameters were maintained within the ranges recommended by Hanlon & Forsythe, 1985, as follows: temperature: 15-20°C; salinity: 34-36 ppt; pH: 7.7-8.2; ammonia: <0.2 mg/L; nitrite: <0.2 mg/L; and nitrate: <100.0 mg/L [36]. A reverse osmosis/deionization filtration system provided automatic top-off water to each tank, and saltwater was made by mixing Tropic Marin Classic Sea Salt with the RO/DI water. More detailed tank and environment set-up and protocols are described by Van Buren et. al., 2021 [19].

All octopus tanks contained at least one den in the form of a small flowerpot or ceramic rock cave, and some octopuses rearranged live rock or other tank items to construct their own dens. Plastic aquarium plants, live rock, and seashells were also frequently used by the octopuses to further conceal dens. Octopuses were either fed live fiddler crabs or frozen-thawed shrimp. Novel enrichment items such as Legos, shells, and other live foods (such live snails, shrimp, and other species of crabs) were provided on a rotating basis to prevent habituation and boredom. Feeding the octopus shrimp in a test tube was introduced as a method of enrichment. To train the octopuses to open the test tube by unscrewing the cap, octopuses were first presented with the test tube containing a piece of shrimp without the cap. If the octopuses removed the shrimp from the uncapped test tube successfully, at the next feeding the cap would be screwed on loosely. As they succeeded in opening the loosely capped tube, the cap would be gradually tightened in subsequent feedings to increase the difficulty of opening it. This process continued with each feeding until the octopuses could remove a cap that was fully screwed on.

To monitor the octopuses, Wyze Cam V2 and V3 cameras were set up facing the front of each tank. The cameras operated 24 hours a day and recorded all motion events to a micro-SD card. In addition, the Wyze phone app allowed for manual recording of live videos to the user’s phone using a start/stop recording function. Videos and pictures taken with portable video cameras were also analyzed.

Burghardt’s [7] five criteria for play were used to determine whether observed behaviors constituted play. Briefly, these criteria are as follows: 1) there is no immediate function to the behavior in the context in which it is performed; 2) the behavior is voluntary, spontaneous, or autotelic (done for its own sake); 3) there is a structural or temporal difference from other typical behavior; 4) the behavior is repeated but not stereotyped; and 5) the animal is healthy and free of stress and competition.

Additional criteria was used to determine the intensity of play behavior based on the “Play Intensity Scale” described by Kuba et al., 2006 [20]. Holding the object close to the mouth was classified as level 0 behavior, exploring the object with the arms as level 1 behavior, and completing one release-grasp sequence (comparative to the push/pull sequence described by Kuba et al., 2006) was classified as level 2 behavior. Level 3 behavior, which was characterized by 2-5 release-grasp sequences, was considered “play-like,” and level 4 “play” behavior consisted of the release-grasp sequence being repeated over 5 times. Release-grasp sequences were only counted when they were performed continually.

## Results

Of the nine octopuses, only three (octopuses E, F, and H) learned to unscrew the test tube cap to access the shrimp within. The characteristics of all nine octopuses are shown in Table 1. Because the test tube cap was not presented to octopuses that had not learned to unscrew it, only these three octopuses had access to the free-floating test tube cap.

**Table 1.** Characteristics of the nine octopuses housed in the lab. F = female; M = male.

| Octopus ID | Sex | Average Weight (g) | Opened Test Tube Cap |
| --- | --- | --- | --- |
| A | F | 229.5 | no |
| B | M | 172.0 | no |
| C | F | 213.0 | no |
| D | F | 153.3 | no |
| E | M | 128.2 | yes |
| F | M | 220.8 | yes |
| G | M | 137.5 | no |
| H | F | 170.5 | yes |
| I | F | 147.8 | no |

**Table 2.** Characteristics of play behavior with test tube caps in octopuses E, F, and H.

| Octopus ID | Number of Times Observed Playing | Skin Patterns During Play | Arms Used |
| --- | --- | --- | --- |
| E | 1 | Acute mottle | L1, L2, R1, R2 |
| F | 5 | Acute mottle | L1, L2, L3, R1, R2, R3 |
| H | 2 | Uniform pale brown phase | L1, R1, R2, R3, R4 |

In accordance with the Play Intensity Scale discussed in the methods, all three octopuses who had access to the free-floating cap engaged in level 4 object play with it. None of the octopuses, including those that were only given the test tube without the cap, chose to play with the test tube itself despite its similar floating properties. For all three octopuses, play consisted of the octopus releasing the cap from its grasp so that it floated upwards into the water current of the tank. As it began to float past, the octopus would reach out to grab it again, pull it in close to their body, then release it again (Figure 1). A video capturing one instance of octopus H playing can be found in the supplemental date (S1 Video).

**Fig 1.**
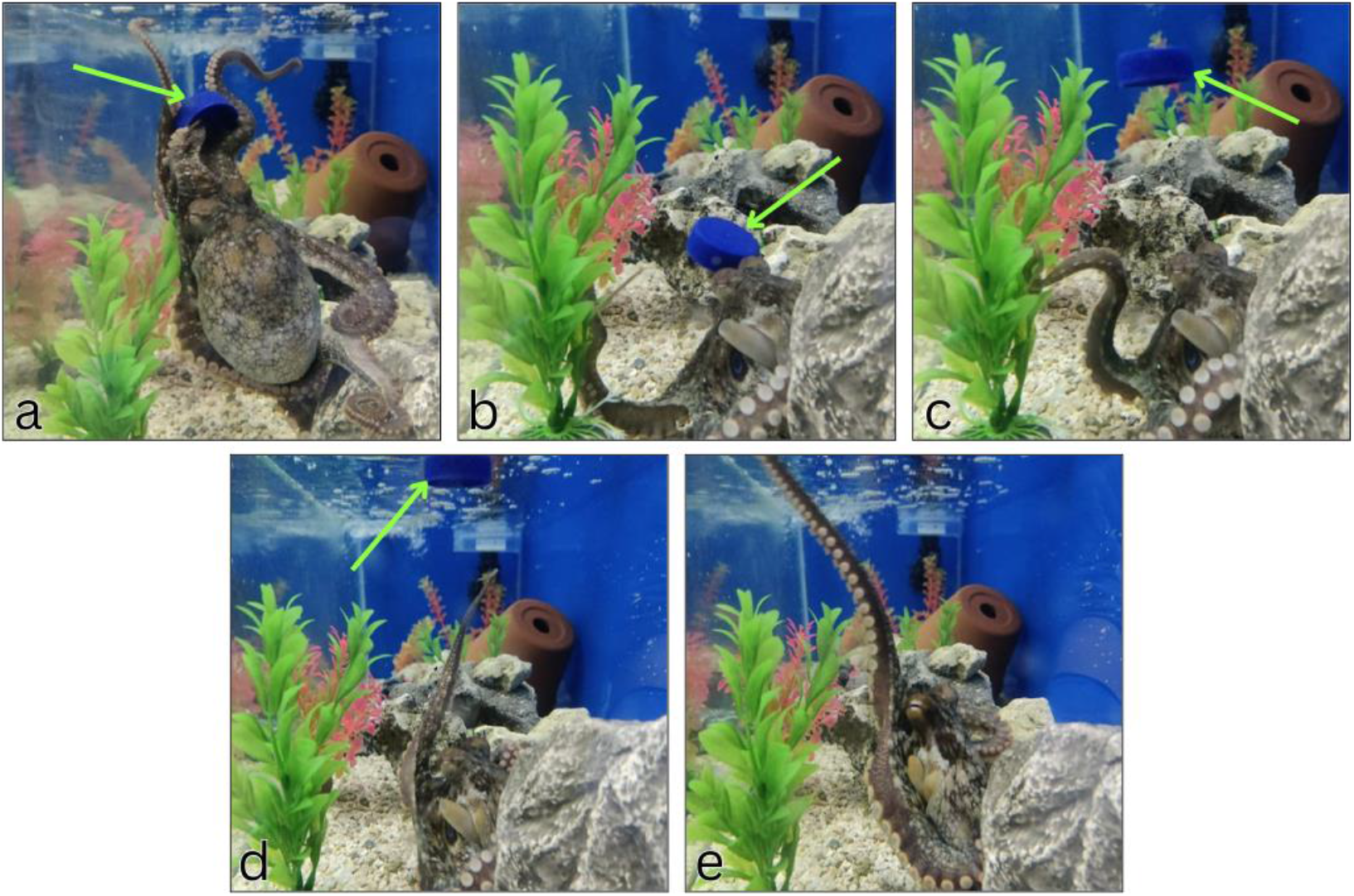
Octopus play sequence with test tube cap. An adult *O. bimaculoides* (Octopus F) grabs a test tube cap with his arms (a), pulls the cap towards his body (b), then releases it into the tank current (c) and reaches to grab it again as it moves away (d, e). The blue cap is indicated by a green arrow in pictures a-d, and is not visible in picture e. This release-grasp sequence was observed in this individual octopus on five separate occasions throughout the four months it was housed in the lab.

The arms used to grasp and release the test tube cap were primarily the front two arms on either side (R1, L1, R2, L2). In some cases, such as when an octopus was using those arms to hold onto the test tube still, other arms were used (R3 and R4). In all octopuses, at least two of the rear arms were used as an anchoring point to a surface in the tank during play, such as the substrate, live rock, or the inside of the den.

## Discussion

This work demonstrates that *O. bimaculoides* engages in play behavior, a behavior that appears to satisfy intellectual and emotional needs. Elucidating this behavior is important to understand the role of play in development and as a tool to assess animals’ wellbeing. These findings also invite further research into how play behavior may vary between octopus species, or into what factors may have contributed to the occurrence of play in individual octopuses, such as life stage, sex, or personality. Mather and Anderson [37] determined that octopuses (*Octopus rubescens*) have individual personalities, characterized by three factors: activity, reactivity, and avoidance. These personality traits may influence the likelihood of an octopus to engage in play activity. The octopuses that displayed play behavior were often active during daytime hours and sought to engage handlers during daily tank maintenance. Comparatively, the octopuses that frequently withdrew from handlers and remained in their dens did not show any signs of play behavior. There is a possibility that the difference in personality between the octopuses housed in the lab could have influenced the likelihood of an individual octopus both to learn how to open the test tube, and to engage in object play with the plastic cap.

In conclusion, the behavior of these *O. bimaculoides* is consistent with previously established criteria for animal play. These findings are also aligned with previously established object play behavior in other octopus species. While there is not enough data here to make any generalized conclusions about the overall correlation between octopus play and welfare, the three animals presented here were determined to be in positive welfare states, and this paper provides preliminary evidence for play behavior in *O. bimaculoides* that may inspire future research. The full characterization of behavior in a species that is increasingly popular in captive settings is an essential aspect to be understood in order to ensure proper care and husbandry methods. A controlled experiment analyzing the play behavior of *O*. bimaculoides, what husbandry factors may impact the occurrence of play, and how the presence or absence thereof correlates with the individual’s welfare state could assist greatly in the development of more detailed and comprehensive standards of care for these animals.

---

**S1 Video. *O. bimaculoides* playing with test tube cap**.

## Notes

### Competing Interest Statement

The authors have declared no competing interest.

### Summary of Updates

Table 1 revised; Table 2 added; Supplemental files updated, Introduction, methods, results and discussion revised for conciseness and clarity.

